# Evolution of object identity information in sensorimotor cortex throughout grasp

**DOI:** 10.1101/2024.10.11.617909

**Authors:** Yuke Yan, Anton R. Sobinov, James M. Goodman, Elizaveta V. Okorokova, Lee E. Miller, Sliman J. Bensmaia

## Abstract

The transition from hand opening to grasping a coffee cup happens smoothly, yet the neural systems underlying this behavior undergo a significant state change at the onset of grasp. At this moment, the motor system begins to control contact forces and the somatosensory system receives a barrage of cutaneous signals that convey information about contact forces and the object’s local features (e.g., edges, texture, and curvature). These cutaneous signals supplement the ongoing flow of proprioceptive input that encodes hand posture as well as muscle forces. How object identity is represented in sensorimotor cortex across this transition remains unknown. In the present study, we sought to quantify the object-specific neural signals in individual neurons of the primary motor cortex (M1) and Brodmann areas 3a, 3b/1, and 2 of the somatosensory cortex in macaque monkeys. Before contact, object-specific information was carried mainly by M1 and proprioceptive area 3a, but this information did not generalize between the periods before and after contact. This observation is consistent with the abruptly changing force signals at contact affecting the assumed postural representation of the object, rather than each modality maintaining an invariant identity. After contact, despite a general decrease in firing rates, information about object identity increased and was encoded with high efficiency across sensorimotor cortex. Cutaneous areas 3b and 1, largely uninformative before contact, became highly informative once objects were grasped. Area 2, which receives both cutaneous and musculotendinous inputs, conveyed little object-specific information before contact, when it too became strongly informative, consistent with its integrative role. Thus, M1 and 3a serve as the main carriers of object information before contact, while cutaneous and integrative somatosensory circuits dominate after contact. This shift highlights the profound change in the coding of object identify in the sensorimotor cortex during contact.

## Introduction

When we reach to grasp an object, we pre-shape our hand to it before making contact ^1–3^, then exert just enough force to grasp and move it without dropping it ^4^. Sensorimotor circuits sense and control the posture of the hand before contact, but they also control force after contact ^5,6^. Lesion studies, leading to manual deficits, clearly implicate the somatosensory and motor cortices in these processes ^7–9^.

Vision allows the hand to be shaped to the object prior to contact, feeding the information about the object shape to the motor cortex ^10–13^. At the same time, information from proprioceptive sensors in the muscles and tendons about the hand posture are conveyed to the motor cortex by the somatosensory cortex ^10,11,14,15^. At the moment of contact, all areas of sensorimotor cortex experience an abrupt change in afferent input about the object. Cutaneous receptors begin to convey information about its local features and pressure on the skin ^16,17^, while musculotendinous receptors report the force exerted by muscles to hold on to the object ^14^. As previous research has largely focused separately on either the pre-contact or post-contact periods, it is not clear how the abrupt transition at contact affects the object-specific signals in the sensorimotor cortex.

In the present study, we characterized the responses of neural populations in sensorimotor cortex after contact with the object was established and compared them to their pre-contact counterparts. We measured the responses of neurons in motor (Brodmann’s area 4) and somatosensory (areas 3a, cutaneous 3b/1, and 2) cortices to understand the evolution of object-related signals distributed across the different cortical fields during grasp. In particular, we investigated the extent to which object-related information is carried by neurons that receive musculotendinous input (in area 3a), neurons that receive cutaneous input (in areas 3b and 1), and neurons that integrate these two streams of information (area 2). Neurons in primary motor cortex (M1) also receive both cutaneous and musculotendinous signals ^15^, which affect its activity during prehension ^18^. Ultimately, we sought to understand how object contact influences the representation of object identity in the sensorimotor cortex.

## Results

We recorded the responses evoked in somatosensory and motor cortices – including Brodmann’s areas 3a (34 neurons), 3b/1 (65 neurons), 2 (36 neurons), and 4 (280 neurons) – from three rhesus macaques as they performed a grasping task (Figure 1A, Supplementary Figure **1**). In brief, a robot presented one of 24 objects that varied in size and shape, directly to the monkey’s hand, eliminating the need for a reach. The monkeys’ proximal arm was restrained to minimize movement. Some of the objects with asymmetric shapes were presented at different orientations, yielding 35 different grasp conditions, each of which is treated as a separate object of different shape or size throughout the analyses. The monkey’s task was to grasp the object with sufficient force to break the magnetic bond to the robotic arm when it retracted. Each condition was repeated six to ten times in each experimental session. As has been previously shown, monkeys (and humans) pre-shape their hand to an object, as evidenced by the fact that object identity can be inferred from hand posture well before contact (Figure 1B, ^10^). Although the timing of the curves was similar, the peak accuracy varied between monkeys, thus, comparisons were made mostly within monkeys. While we have previously characterized the neuronal responses in this task before object contact^10,11^, the objective of the present study was to compare this representation to that after contact, when neural activity reflects not only the posture and movement of the hand but also contact with the object.

**Figure 1.**
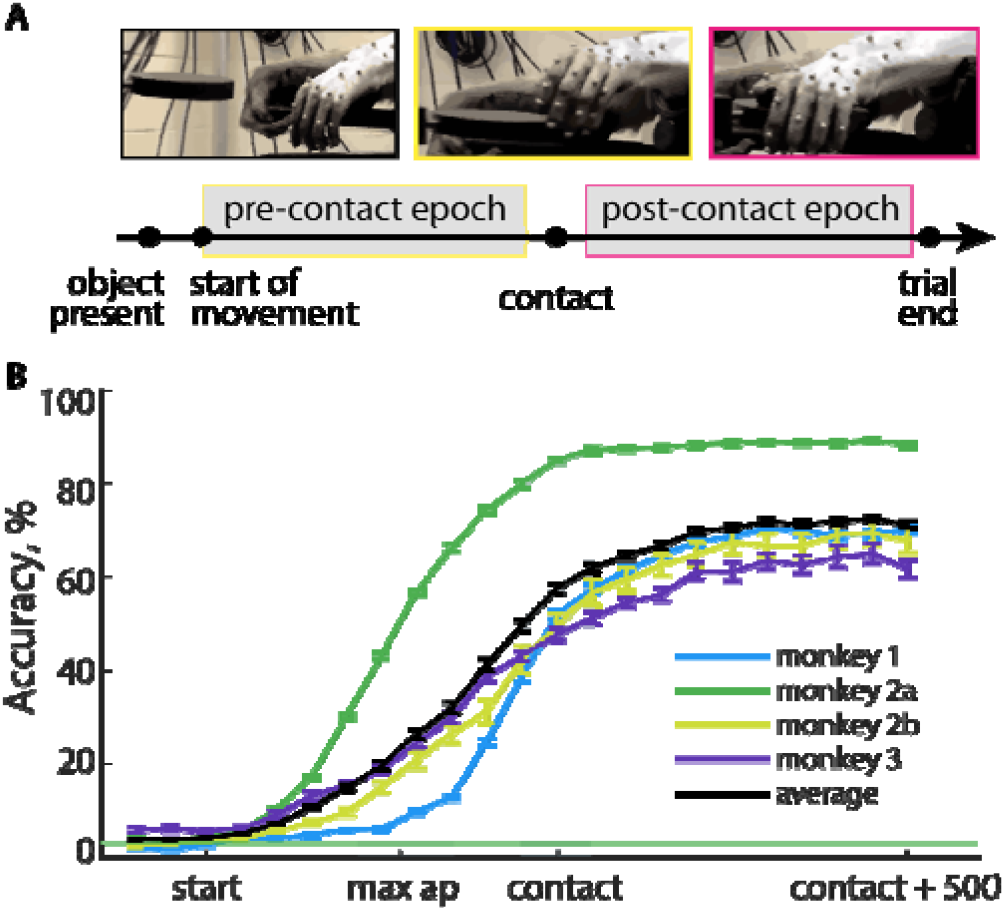
Behavioral task. **A**| Top: images of one of the trials, taken during object presentation, before, and after the contact. Bottom: Task schematic. The analysis has focused on the pre- and post-contact epochs. **B**| Object classification based on hand kinematics (joint angles). Classification accuracy is computed as a function of the time from 100 ms before the start of movement to 500 ms after contact. Accuracy is averaged over three monkeys across all sessions. Error bars denote SEM. The chance level (2.9%) is shown in green. Dark grey shaded area denotes peri-contact period

We identified individual neurons across all recorded areas that were modulated by the task and conditions (Figure 2, Methods Table 1). Neurons in M1 and area 3a often showed changes in firing rate prior to contact with an object, whereas neurons receiving cutaneous inputs (areas 3b, 1, 2) exhibited strong phasic responses at the moment of contact (Figure 2AB). To quantify the typical responses and the timing of object information in sensorimotor cortex, we examined the activity of individual neurons and their relation to the object identity.

**Table 1.**
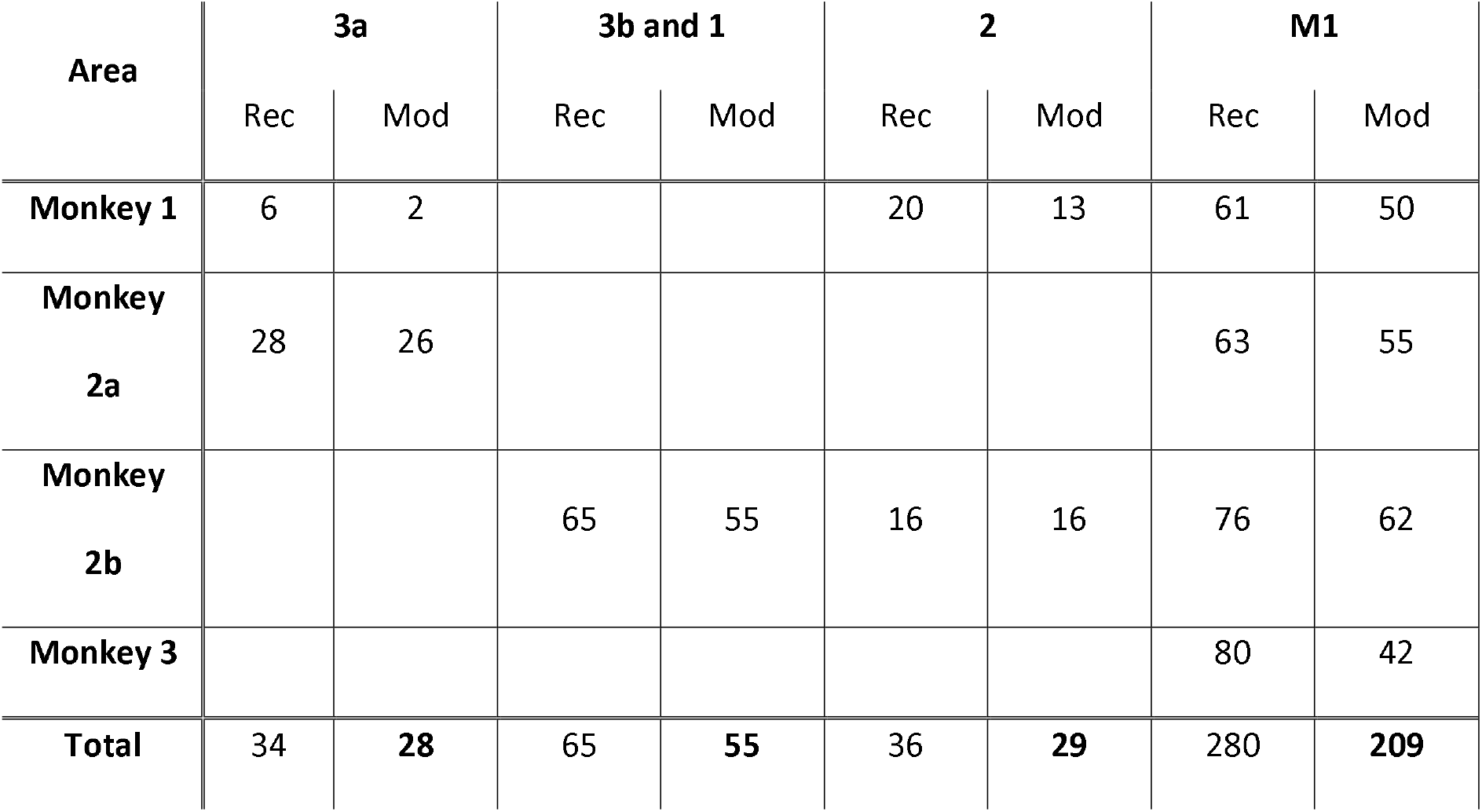
Distribution of recorded and modulated units between monkeys. Rec: recorded; Mod: modulated to the task and conditions.

**Figure 2.**
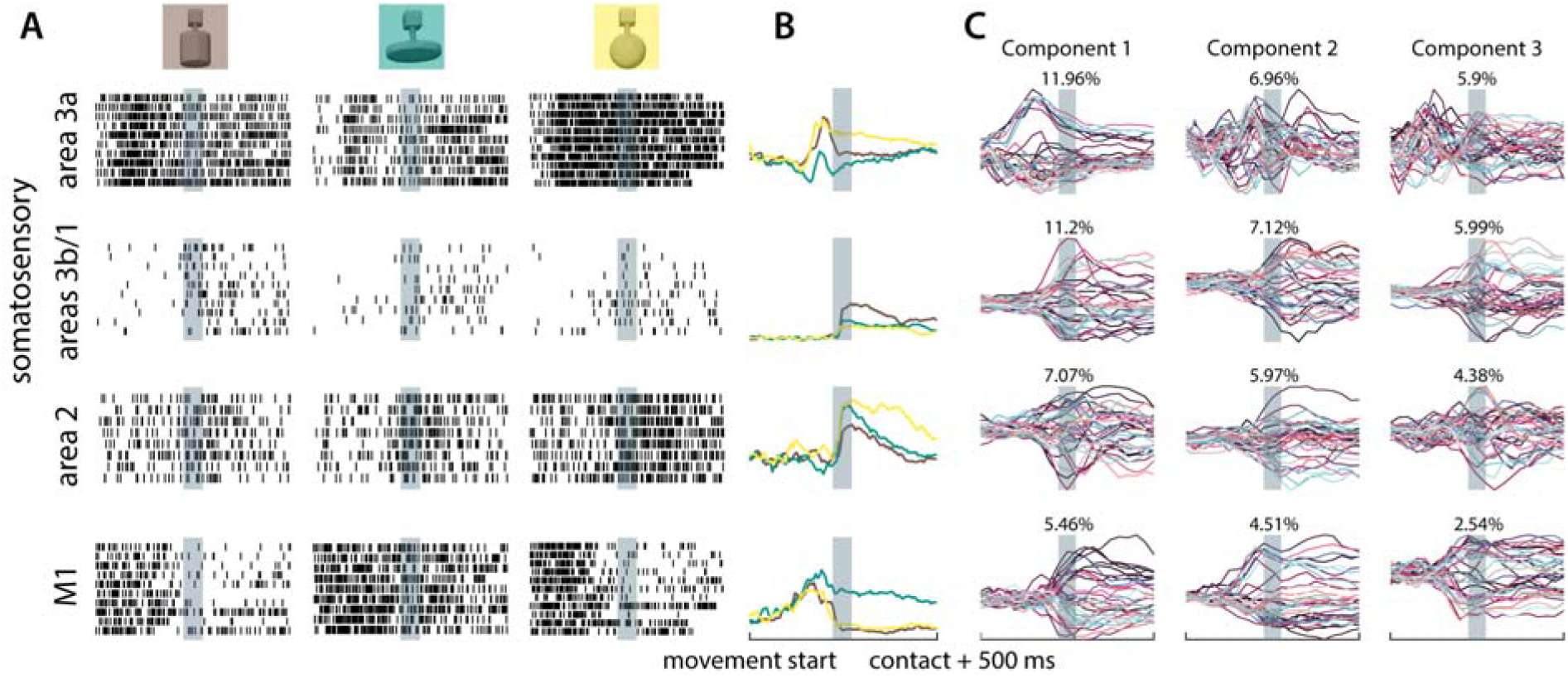
Activity of neurons in sensorimotor cortex during grasp of individual objects. **A**| Raster plots of one neuron from each somatosensory and motor area as the animal grasped 3 objects over an approximately 1200-ms window centered on object contact. The period before contact has been warped in time from the beginning of the movement to the estimated time of contact between the fingers and the object. The shaded regions show the peri-contact epoch, the estimated contact was in the middle of that period. **B**| Average firing rates of the neurons from A. Colors denote the objects shown in A. **C**| First three object identity-dependent components of neural activity calculated using dPCA. Each color represents neural activity averaged during the monkey’s interaction with one of the 35 objects, after projection onto the component specified by the column. The number indicates the amount of variance explained.

Across areas, the condition-dependent components emerged and decayed at different times, highlighting the distinct temporal profiles of object-related encoding in each region. To visualize these dynamics, we performed demixed principal component analysis (dPCA)^30^ for each area (Figure 2C). We projected the condition-averaged neural activity onto the first three condition-dependent components. We focused on the condition-dependent components, since the condition-independent component was closely related to the average activity in each region, which we consider later (Figure 3). The separation of these trajectories confirmed the presence of information about the object identity in neuronal populations. In M1, trajectories diverged prior to object contact, consistent with its role in movement planning based on the visual inputs. Similarly, but presumably due to proprioceptive input rather than motor planning, trajectories in area 3a also separated prior to contact. The almost categorical separation of object traces in the first 3a component was driven largely by a corresponding difference in the wrist angle as the two groups of objects were grasped. In contrast, the trajectories in areas 3b and 1, as well as in area 2, diverged around the time of contact and remained distinct after the object has been grasped, consistent with their somatosensory-driven response properties.

**Figure 3.**
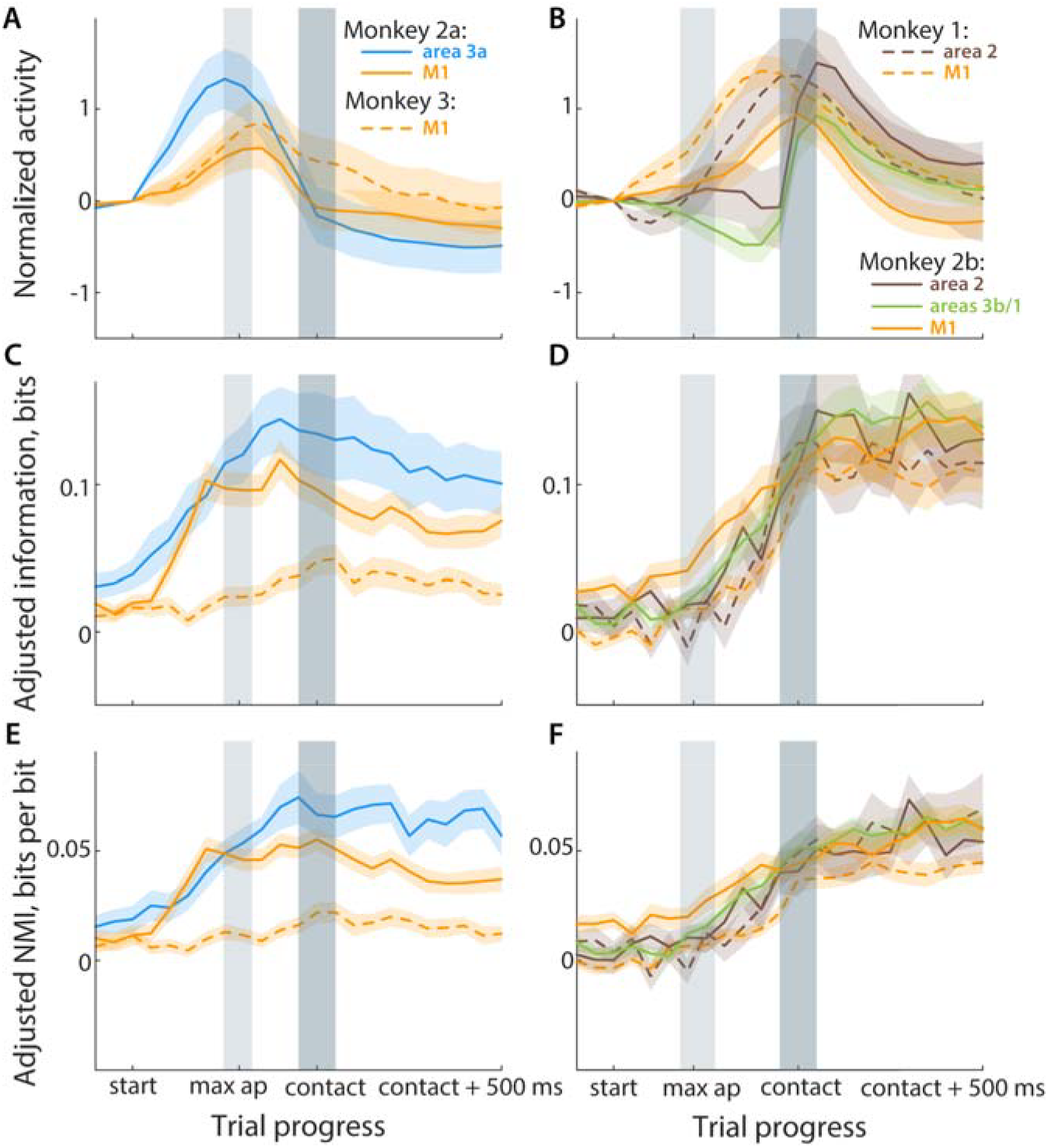
Time course of activity of individual neurons and their informativeness in sensorimotor cortex. **AB**| Average normalized firing rates across all neurons pooled across monkeys for each of the somatosensory areas and M1. Normalized firing rate is calculated by subtracting the raw firing rate at the start of the movement and then dividing by the standard deviation. **CD**| Mutual information between the activity of a neuron in each area and object identity as a function of time, adjusted to chance level at each time point. **EF**| Same as C and D, but the mutual information was normalized to neuron’s entropy before subtracting chance NMI. Max ap: Maximum aperture, NMI: Normalized mutual information. Light gray region denotes interquartile region of maximum aperture moment (43-71% of the period before contact, median=59%), dark grey – peri-contact period. Shaded region denotes SEM.

### Population activity in sensorimotor cortex during grasp

We examined the evolution of the population activity as the monkeys grasped each object, beginning 100 ms before movement onset through 500 ms after object contact (Figure 3AB). In area 3a, activity peaked when the hand reached its maximum aperture before grasp (Figure 3A). After contact, however, responses in area 3a were weak, even though the monkey was exerting force on the object. This weaker activity during grasp was unexpected, as proprioceptive neurons in 3a have been shown to encode not only kinematics but also force ^5,19^. Note, however, that muscle activity also typically drops after the initial grasp ^20^, which may be the cause of decreased afferent input.

In areas 3b and 1 the activity was weak during the reach but surged after contact, as expected, given that neurons in these cortical fields have exclusively cutaneous inputs (Figure 3B). That responses were slightly higher at reach onset than later in the pre-shaping epoch likely reflects cutaneous responses through contact with the armrest. Area 2 activity increased before contact in monkey 1, but not in monkey 2, where the responses followed the activity profile of areas 3b/1 more closely, although without the dip in activity before contact. This pattern is consistent with the combined proprioceptive and cutaneous input to area 2 ^21–25^.

Finally, in M1, responses rose during the pre-shaping, peaked somewhat later than 3a, then declined after contact. As in area 3a, the decrease in M1 activity is surprising, given that the monkey was exerting substantial force on the object during this epoch. It might reflect control taken over by subcortical structures and spinal cord ^26,27^.

Having established the aggregate neuronal response before and after grasp in each cortical field, we next examined the time course of the object-specific information over the same period (Figure 3CD). To that end, we computed the mutual information ^28,29^ between the activity of each neuron and the identity of the grasped object in a 160 ms sliding window throughout the trial. Trial-shuffled information was subtracted to correct for chance correlations. Broadly speaking, the information followed the time course of neural activity, with several notable exceptions. Cutaneous areas 3b and 1, as well as area 2 experienced an increase in information after contact (Figure 3D, p<0.001, r=-0.77 and r=-0.67, Supplementary Figure 2), while proprioceptive area 3a – a weak decrease (Figure 3C, p=0.37, r=0.18).

Unexpectedly, the information in the individual neurons of area 2, despite its combination of cutaneous and proprioceptive inputs and relatively elevated activity before contact, did not rise above that of the purely cutaneous areas 3b and 1. It is possible that the input from cutaneous areas impedes area 2 from forming a meaningful representation of the object identity based on proprioceptive inputs. Although M1 activity rose before that of area 2 in monkey 2b, the informativeness of both of these signals increased much later in monkey 1. Furthermore, while M1 began to rise a bit earlier, area 2 reached its peak informativeness before M1 did. Overall, M1 neurons trended toward increased information after contact, although the difference was only significant in monkey 2 (p=0.003, r=-0.21; Supplementary Figure 2). Comparable information levels before and after contact likely reflects the early rise in M1 information. It most likely reflects the input from the visual system into M1, which guides the formation of object-specific kinematics. Interestingly, the decrease in activity after contact in each area was not reflected in the information that activity represented.

To test whether the decline in activity reflects an increase in the efficiency of information encoding, we computed normalized mutual information (NMI) by dividing the information of each neuron by its entropy (Figure 3EF). Higher NMI would indicate that more of neuron’s activity is informative about the object being grasped. As with information, we have adjusted to the chance level of NMI at each time point. In all areas, NMI plateaued around the moment of contact (p<0.003, r=-0.59, r=-0.56, r=-0.23 for areas 3b/1, 2, and M1, respectively) then remained stable, except for area 3a, whose NMI was similar before and after contact (p=0.25, r=-0.23). In Monkey 3 M1 the effect was not significant (p=0.29, Supplementary Figure 3). This suggests that the object information was consolidated and preserved through the decrease in overall activity.

### Object-specific signals in neural populations

The mutual information analyses gauge the degree to which individual neurons in each cortical field carry object-specific information. To compare the overall informativeness of each area about the objects, we analyzed the degree to which object identity could be inferred from the activity of equal numbers of random subgroups of neurons (Figure 4). This analysis was performed on monkey 2 data, where recordings were available from all four areas. Furthermore, we excluded ±50 ms around the contact from the analysis to avoid misattributing the precisely-timed, phasic signals from neurons with receptive fields with very early object contact to the precontact period (Figure 2AB,^17^). By excluding this period, we could focus on stable states before and after contact.

**Figure 4.**
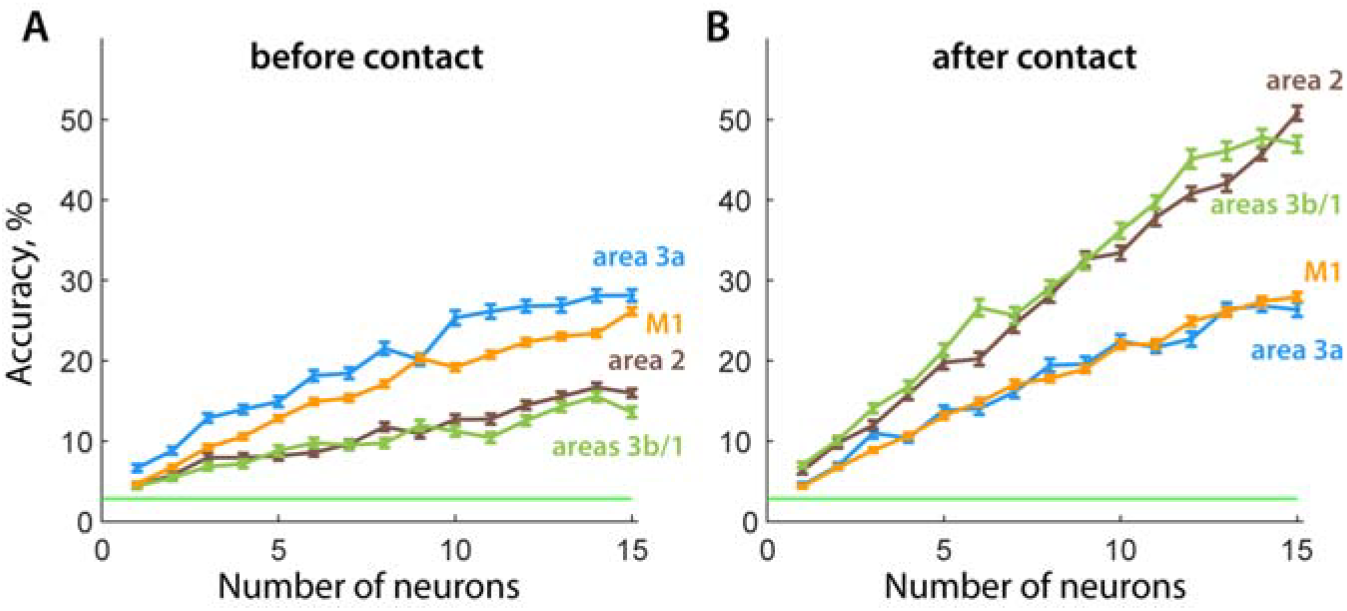
Classification of objects based on population responses. **A**| Performance of a classifier of object identity based on the responses of neurons in sensorimotor cortex before contact, as a function of sample size. **B**| Classification accuracy after contact. Green line denotes chance. Error bars denote SEM.

We found that the cortical areas differed in their performance before and after contact (Kruskal-Wallis test p<0.001). Before contact, M1 and area 3a had a similarly high performance, surpassing somatosensory areas 3b, 1, and 2 (Mann-Whitney U-test p<0.001, rank-biserial correlation r=0.72). They maintained the high performance after the contact, while areas 3b, 1, and 2 surged to almost twofold population-level representation of object identity (p<0.001, r=0.9). Notably, the strength of the representation of object identity in area 2 population was comparable to that of areas 3b and 1 (p=0.003, r=0.25). This outcome was unexpected, given that area 2 also receives proprioceptive information from area 3a – a feature that would have predicted a stronger object-related signal than in 3b/1 before and after contact.

We have demonstrated that all the areas we recorded from carry object-specific information. However, the outstanding question is whether that information is encoded similarly before and after contact, i.e., does it persist through the establishment of contact. To assess the stability of the representation, we repeated the analysis using classifiers trained on data from one epoch and tested on the other (Figure 5). We reasoned that a purely kinematic representation would be stable across epochs because hand posture immediately before contact is similar to that just after. Instead, we found that the cross-epoch classifiers (Figure 5, purple) all performed poorly, indicating that the object representations changed dramatically upon object contact. In areas 3b, 1, and 2 they performed at chance level and in M1 and area 3a their performance decreased significantly (Mann-Whitney tests, p<0.001, r>0.93).

**Figure 5.**
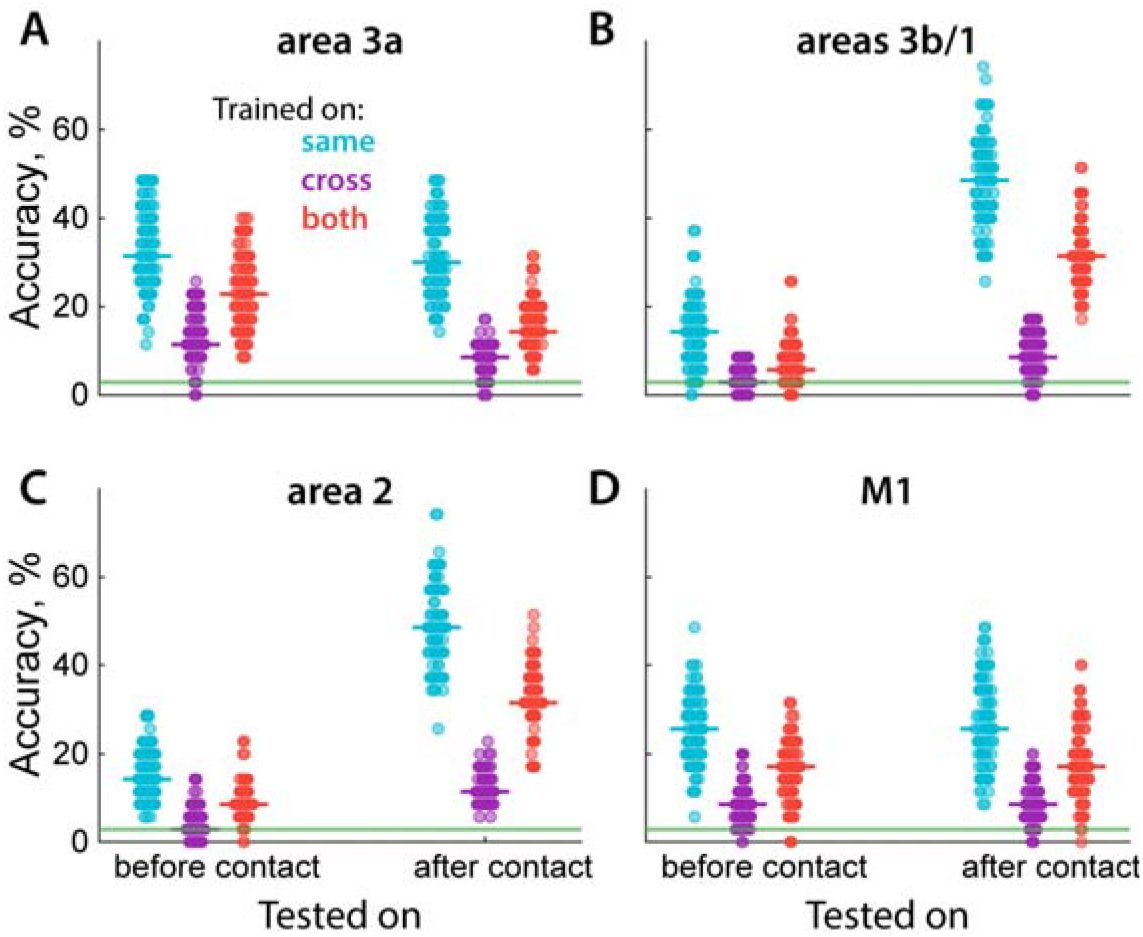
Stability of object-specific signals across epochs. **A**| Performance of classifiers of object identity based on the responses of neurons in area 3a trained and tested within the same epoch (cyan), trained in one epoch and tested in the other (purple), and trained in both epochs then tested in each (red). The “before contact” vs “after contact” column indicates the test epoch. Each point denotes the accuracy of classification based on a random allocation of training and testing data and the bold line denotes the median performance. **B, C, D**| Same as A for areas 3b/1, area 2, and M1, respectively. Horizontal green line denotes chance (1/35).

Although the decoders did not generalize well with training data limited to only one epoch, we hypothesized that there might be a unified representation preserved throughout the behavior. To test this, we trained a classifier on combined pre- and post-contact data and evaluated its performance for each epoch separately (Figure 5, red). Even with this more robust training data, in somatosensory areas 3b/1 and 2, the classifier performed poorly on the pre-contact epoch, corresponding to the lack of postural information prior to contact in areas 3b/1 and suggesting a strong effect of cutaneous information on the potentially informative proprioceptive information in area 2. Although this classifier performed better than chance in both areas 3a and M1, it still suffered a significant drop in performance from classifiers trained and tested on a single epoch (median drop in accuracy 40% and 34%, p<0.001, r=0.69 and r=0.57, respectively), consistent with the interpretation that these two areas carry information about hand posture. The drop in the decoders’ performance suggests that signals related to contact forces are integrated with the postural signals in these areas. Interestingly, the drop in performance was more pronounced after contact than before in area 3a (26% drop vs 53% relative to within-epoch classifier), implying that postural information is integrated with cutaneous after contact, and thus impacts the postulated purely postural representation. Similar results were observed for monkeys 1 and 3 (Supplementary Figure 4); and were further supported by joint angle decoding from the neuronal population, which exhibited trends consistent with these results (Supplementary Figure 5). Collectively, these findings indicate that all areas hold information about the grasped object following contact, and those representations are affected by the forces exerted on the grasped object.

## Discussion

Manual dexterity is facilitated by an expansive neural network that spans large swaths of the cortex ^31–33^. Different nodes within this network are thought to make different, albeit overlapping contributions to hand control, so it is not surprising that contact with an object would affect the sensorimotor areas in distinct ways (cf. ^13^).

In preparation for grasping, the hand pre-shapes to match the object, allowing for an efficient and effective grasp. It also allows object identity to be inferred from the shape of the hand with high accuracy, even before the hand touches the object ^34,35^. The proprioceptive information about limb posture and movement transduced by muscle receptors (muscle spindles and Golgi tendon organs) is received by area 3a of the somatosensory cortex ^5,21,36–40^. It is not surprising then that the activity of area 3a peaks before contact, as the hand reaches its maximum aperture ^10,11^. Interestingly, while the response strength peaks at maximum aperture, its informativeness peaks closer to the time of contact, as the hand begins to conform to the object. Activity in 3a drops sharply after contact, although surprisingly, without a significant impact on the informativeness of individual 3a neurons. A similar phenomenon of decreased activity during contact has been previously documented in a subgroup of cortical somatosensory neurons ^13,41^. Furthermore, muscle activity also decreases after contact ^20^, which might reduce the signal coming from proprioceptive sensors.

We did not see strong object-specific information in M1 at the start of the movement, when visual inputs from parieto-frontal networks are typically thought to guide motor planning^13,32,41^. Instead, object identity signals in M1 peaked later, together with those of proprioceptive area 3a. It did not, however, affect the macaques’ ability to preshape their hands in preparation for grasping the object. It should be noted that 3a neurons were recorded only in one monkey. After contact, M1 activity declines, which conceivably drives the aforementioned decrease in muscle activity. From a functional perspective, the control of the hand by M1 shifts to force maintenance in an isometric posture. The addition of the force control signal to M1 activity arguably makes it even more object-specific. However, the informativeness of M1 neurons after contact was similar to that before contact (Figure 4), and lower than that of the somatosensory areas, perhaps a product of the decreased activity levels in M1 after contact. The decrease in activity might reflect the control of hand motion being passed down to subcortical structures and the spinal cord. For example, reticular formation has been implicated in the control of gross hand movements ^27,42,43^, and magnocellular red nucleus in monkeys^44–46^. The spinal interneurons, in turn, often exhibit sustained responses during isometric wrist tasks as opposed to a phasic encoding of variables in the motor cortex ^26,47^. Although the contribution of the cortex to control might decrease, our results suggest that it still holds object information throughout the task.

While the information about the object is always present, it changes after contact, highlighting the entanglement of postural and kinetic signals. If the signals were fully kinematic and unaffected by force, decoders built on pre-contact epochs or on both epochs might be expected to perform as well on post-contact epochs as on their original domain, but they do not. While these points present a different view from the previously proposed orthogonality of postural and force signals in M1 ^48,49^, we cannot test it directly because the experimental setup did not record interaction forces or electromyograms. Furthermore, that the decoders built on data from both epochs combined can extract some postural representation, which is invariant to the contact.

In stark contrast to areas 3a and 4, areas 3b and 1 are nearly quiescent before contact, with weak signals that are almost completely uninformative about the object. This response pattern aligns with the fact that areas 3b and 1 receive exclusively cutaneous input ^25^, suggesting that pre-contact cutaneous signals are largely uninformative about hand posture. While cutaneous signals have been shown to convey information about the posture of individual joints ^50,51^, our results suggest that they do not encode hand state under naturalistic conditions, where multiple joints are deflected simultaneously ^11^. Areas 3b/1, however, become highly informative about the object after contact. Following contact, there are at least two ways in which cutaneous signals might be informative about an object in the current experimental paradigm, where the object is handed to the animal in a specific way. First, the pattern of contacts on the skin might vary across objects, which would be reflected in different patterns of activation across the neural population ^25^. Second, cutaneous signals might carry information about the local features of the object – such as edge, curvature, and texture – at each point of contact. Both of these sources of cutaneous information could contribute to the performance of object decoders based on activity in areas 3b/1 after contact.

Area 2 exhibits response properties that have a lot of similarity with areas 3b and 1. Indeed, the strength of the response in area 2 is higher than in 3b and 1 throughout grasping but informativeness about objects keeps rising before the moment of contact and is maintained at a high level after contact, similar to purely cutaneous areas 3b and 1. Although area is the first stage along the somatosensory neuraxis where signals from the skin and signals from the muscles and tendons are integrated ^21–23,52^, information about object identity or ability to classify the grasped object using populations of neurons does not rise above those of areas 3b and 1. It is additionally surprising that area 2 is relatively uninformative about objects before contact. One possibility is that information about hand posture inherited from area 3a is obscured by cutaneous inputs inherited from areas 3b and 1. Those responses reflect the skin stretch during pre-shaping and are not informative about the object’s identity. Then, the neurons in area 2 combine object-specific proprioceptive information with uninformative cutaneous signals, leading to low classification rates before contact. The strong effect of cutaneous signals on object representation is also highlighted by the inability of decoders to generalize between epochs. This highlights the complex integration of somatosensory information in area 2 that guides object and shape recognition.

Although this study was somewhat limited by the number of neurons recorded in each area, key findings related to the multimodal areas 2 and M1 were replicated across two or three monkeys. Results from the predominantly unimodal areas 3a, 3b, and 1 were consistent with the previously published literature. A further limitation is the absence of contact force recordings, which limits our ability to precisely identify pinpoint the timing of contact events and to assess the influence of grasp force on sensorimotor cortical activity. Incorporating force sensors in future experiments would allow for a more nuanced interpretation of neural correlates of force production during grasp. Although previous work has shown that wrist movement carries information about object identity^53^, we did not investigate commonalities across object orientations in this study. This decision was due to task constraints that prevented the monkeys from fully rotating their wrists, effectively resulting in different grasping strategies, i.e., grasped object, for differently oriented presentations. It remains possible that M1 and area 3a contain a common non-linear neural subspace containing information of the grasped object that is fully preserved through contact, however, our attempts with several non-linear methods did not reveal such a representation. Despite these limitations, the key findings of this study remain well-supported by the data and align with previous literature. These results lay a solid foundation for future research aimed at uncovering the neural mechanisms of sensorimotor cortex that support dexterous manual behaviors.

## Conclusions

In conclusion, each area of the sensorimotor cortex exhibits distinct patterns of activity during grasping, highlighting the complexity of the neural basis of hand control and somatosensation. Motor and proprioceptive areas are highly informative about object identity already before contact and remain so after contact, though their object representations change. This shift might reflect the musculoskeletal basis of object-related signals, which intertwine the kinetic and kinematic representations. Neurons in areas 3b and 1 are uninformative about object identity before contact, indicating that whole-hand posture is poorly encoded in cutaneous neurons. Area 2 integrates these inputs, becoming highly informative after contact, consistent with its putative role in stereognosis. The weak informativeness of area 2 before contact suggests a strong entangling of cutaneous and proprioceptive signals, obscuring the object representation coming from area 3a. Overall, the effects of contact on the object-related information in the areas of the sensorimotor cortex emphasize their function and anatomy.

## Methods

### Animals and surgery

Three male rhesus macaque subjects from 6 to 15 years old, weighing between 8 and 11 kg participated in this study. All animal procedures were performed in accordance with the rules and regulations of the University of Chicago Animal Care and Use Committee. Monkeys received care from a full-time husbandry staff, and a full-time veterinary staff monitored animals’ health on a daily basis. All monkeys were implanted with a head-fixing post onto the skull. Monkey 1 was implanted with semi-chronic array with electrodes of adjustable depth (Gray Matter Research; Bozeman, MT). Monkey 2 was implanted twice, first with semi-chronic Gray Matter drive, and then with two Utah electrode arrays (UEAs; Black-rock Microsystems, Inc. Salt Lake City, UT). Monkey 3 was implanted with 4 floating microelectrode arrays (FMAs, Microprobes for life science; Gaithersburg, MD) and 2 UEAs. All procedures were performed under aseptic conditions and anesthesia induced with ketamine HCl (20 mg kg^−1^, IM) and maintained with isoflurane (10–25 mg kg^−1^ per hour, inhaled).

### Behavioral task

Monkeys were trained to grasp 35 objects varying in size, shape, and orientation. In a typical grasp trial, the monkey sat in a chair with its head fixed and arms resting on cushioned armrests. The arm was secured near mid-humerus in a circular plastic support that reduced shoulder mobility, ensuring that the arm of the monkey remained largely immobile during the grasp. At the start of the trial, a robotic arm directly facing the monkey presented an object. The monkey was trained to grasp the object and apply enough grip force such that, when the robotic arm retracted, the object would be disengaged from its magnetic coupling with the robot and remain in the monkey’s hand. The stationarity of monkey’s arm was enforced by a photosensor on the armrest which ended the trial (without reward) if the arm was lifted to expose it to light. The period from the grasp to robot retraction was 1 or 3 seconds randomly chosen for each trial. For more details, refer to ^10,11^.

### Electrophysiology

In monkey 1, we recorded from depth-adjustable electrode arrays placed over the central sulcus (Supplementary Figure 1). For monkey 2, we first recorded from depth-adjustable electrode arrays over the central sulcus (first implant), then from the UEAs in the pre- and post-central gyri (second implant). The monkey was implanted for the second time three years after the first. For monkey 3, we recorded from UEAs implanted in the pre-central and post-central gyri and FMAs implanted in the posterior and anterior banks of the central sulcus. Offline spike sorting (Offline Sorter, Plexon, Dallas, TX) was used to isolate individual units and identify spikes.

Somatosensory units were classified based on anatomical locations and manual functional mapping prior to recordings. To differentiate cutaneous, proprioceptive, and combined response types, we functionally characterized each somatosensory cortical unit using a standardized manual mapping procedure (see ^10,11^ for more details). Light tactile stimulation was applied to the hand and arm using a cotton swab to identify cutaneous responses. Proprioceptive responses were assessed by manually manipulating the joints of the hand and wrist, and by palpating forearm muscles and tendons. For neurons responsive to tactile stimulation, we varied wrist and hand positions (e.g., flexion, extension, pronation, supination) to determine whether the response was posture-dependent, indicating a combined modality. Only neurons that responded selectively to stimulation of the upper limb were included in the analysis. Histological mapping has been performed on Monkey 1 to confirm the recording sites (Figure 1E in ^10^). The Monkey 1 corresponds to Monkey 4 in Goodman et al., 2 to Monkey 2, 3 – to Monkey 1. The locations of electrode insertions can be seen in Supplementary Figure 1.

The standard deviation threshold for spike sorting was 5 (matching the default in the Blackrock recording software) and adjusted manually (usually upward) on a case-by-case basis. The likelihood of resampling a given neuron across sessions was low, as most Gray Matter electrodes were repositioned between recording sessions. The recording sessions happened several days apart, which would lead to low probability of recording the same neuron with FMAs and UEAs. In total, we recorded 20 sessions: 9 with Monkey 1, 9 with Monkey 2, 2 with Monkey 3.

From the recorded units we selected those which were modulated by the task and by experimental conditions (Table 1). Task-related modulation was assessed using Friedman paired test on three epochs: baseline (750 ms before movement to movement start); 450 ms before contact to 50 ms before contact; 50 ms after contact to 450 ms after contact. Dependance on condition was assessed using two Kruskal-Wallis tests on the before- and after-contact periods respectively, which were combined into one p-value using Fisher’s method. The threshold p-values were adjusted for multiple comparisons using Benjamini and Hochberg method.

### Kinematics

We measured hand kinematics using a marker-based motion tracking system (Vantage, VICON; Los Angeles, CA). In brief, 30 photoreflective markers were placed on the back of the monkey’s hand, covering major joints of the hand and wrist. Ten to fourteen infrared cameras, placed around the workspace, tracked the markers at 100 Hz for monkeys 1 and 2 and 250 Hz for monkey 3. We then used the Vicon Nexus software to reconstruct the 3D position of each marker, manually labelling markers that were missing due to occlusion. Then, we used a musculoskeletal model (scaled to the monkey’s size) in OpenSim ^54^ to perform inverse kinematics, which inferred the time-varying joint angles of the hand from the 3D marker positions. Joint angles were smoothed with a 50-ms moving average and the angular velocities were computed from these.

### Parsing different trial epochs

For each trial, we identified the start of movement (finger and wrist position deviated from rest) and contact with the object (finger and palm stop moving after conforming to the object). For monkey 1, the events were labelled manually for all trials. For monkeys 2 and 3, we first manually labelled a subset of the trials, then trained a multi-class linear discriminant analysis classifier (LDA) using the joint angular velocities 200 ms before and after each video frame. We used the trained classifier to identify the key kinematic events in the remaining trials. Among all sessions from monkey 2, the mean and the standard error of the model deviation from left-out hand-labeled events were 16 ± 9.5 ms for start of finger movement, 21 ± 8.8 ms for start of wrist movement, –25 ± 9.7 ms for palm contact with an object, and – 48 ± 12.1 ms for finger contact with an object. For more details, refer to ^10,11^.

### Neural data preprocessing

After sorting, neural responses were first divided into non-overlapping 10-ms bins. For each trial, we aligned neural activity with the identified kinematic events, restricting our analysis to the period from 200 ms before movement onset to 500 ms after first contact with the object. The period before contact began at the hand movement onset and ended 50 ms before first contact. As that period varied considerably (IQR: 600-910 ms, 90 percentile: 490-1950 ms, median 720 ms), we resampled it into 10 equally spaced points for each trial. The period after contact began 50 ms after contact and ended 500 ms after contact, and was also divided into 10 equal periods. These 20 timepoints were aligned across trials, and used to compute mutual information and normalized mutual information (NMI) by averaging activity over a 160-ms window centered on each time point. The median period of maximum aperture was at 59% of the before-contact period (IQR 43-71%).

### Mutual information analysis

To quantify the information about the grasped object carried by each neuron’s response, we used both the mutual information and normalized mutual information (NMI), which were computed as follows:

First, we computed the entropy (H) of the neuron’s response:

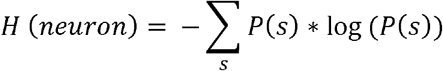

where ***s*** denotes the neuron’s spike count over a specified time window (e.g., 200 ms before contact), P(s) – discrete probability of a firing rate in the specified time window. Then we computed the condition entropy of the neuron, given each object, **o**:

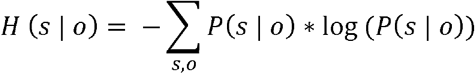

where P(s|o) is the discrete conditional distribution of a firing rate of the neuron for object **o**. Next, we computed the mutual information:

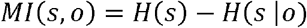

The mutual information measures in bits, the amount of information about the object identity contained in the neural response of a single neuron. We then normalized the mutual information by the neuron’s overall entropy:

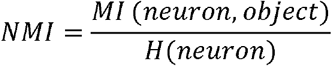

The resulting metric, NMI, ranges from 0 to 1, with 0 meaning that all neuron’s activity is unrelated to the object identity, and 1 – that all of its activity is encoding object identity. We did not include *H* (*object*) in the normalization because this term depends only on the number of trials for each object and was therefore constant for all neurons. Furthermore, normalization to the entropy of the neuron reveals how much of its activity is related to the object identity.

To remove the sporadic correlations from both metrics, we computed chance information and NMI for each neuron at each time point by permuting the object identity on each trial 100 times. We then subtracted the mean of chance distribution from the information and NMI.

### Statistical analysis

To calculate the effect size of the difference in distributions, we used Rank-Biserial correlation (r) for Wilcoxon sign-rank test by dividing the Z-statistic of the test by the square root of the number of observations (Figure 3, Supplementary Figure 2, Supplementary Figure 3). When comparing the unpaired data, we used a Mann-Whitney U test with Rank-Biserial correlation estimate of the effect size r (Figure 4). To test if more than two samples came from the same distribution (Figure 4, Figure 5), we used Kruskal-Wallis test.

### Demixed principal component analysis

We used demixed principal component analysis (dPCA)^30^ to visualize object-specific population activity across 35 conditions. For each animal, single-trial neural responses were aligned as described above and pooled across sessions within each recorded brain area. dPCA was applied separately to each area to extract 20 demixed principal components (dPCs) that captured both condition-independent (not shown) and object-specific variance. We visualized the object-related population dynamics by plotting the time course of neural responses projected onto the top three object-specific dPCs (ranked by percentage of variance explained) averaged across trials for the 35 conditions.

### Classification

To investigate the object information carried by populations of neurons, we used a classification analysis. In brief, we pooled trials across sessions and calculated the spike count for windows before and after contact. Each window excluded a 100-ms buffer zone around contact, for reasons described above. In preliminary analyses, we found that 400 ms windows provided the best classification accuracy across cortical fields. We then used an object-level, leave-one-out classification paradigm: For each object, we randomly selected a test trial and trained a linear discriminant analysis classifier (LDA) on the remaining trials. We calculated the classification accuracy on the left-out test trials. We repeated this process 100 times for each cortical region and calculated the mean accuracy. To assess the effect of population size, we randomly selected subsets of neurons in each repetition and computed the mean accuracy as a function of sample size.

We also performed an across-epoch classification analysis to understand the stability of object signals before and after contact. To this end, we used the same pre-and post-contact windows and leave-one-out paradigm, but tested classifiers trained with data from one window on data from the other. We also trained the LDA classifier on the combined training data from both epochs and tested it on both epochs, separately (“both” classifier in Figure 5 A-D).

## Author contributions

YY, ARS, and SJB designed the study. JMG and EVO performed the experiments. YY, EVO, and ARS performed the analysis. ARS, LEM, and SJB supervised the work at different stages. YY, ARS, and LEM wrote the manuscript with input from all authors.

## Acknowledgements

The authors would like to thank Dr. Stephanie Palmer of the University of Chicago for helpful discussions. The authors would like to thank the reviewers for the interest and suggestions.

## Data availability

All data generated or analyzed during this study as well as analysis and plotting scripts are available in a public repository https://uchicago.box.com/s/ce8z1cofq5jlmolb5uuc51exfx9slen4.

---

**Supplementary Figure 1.**
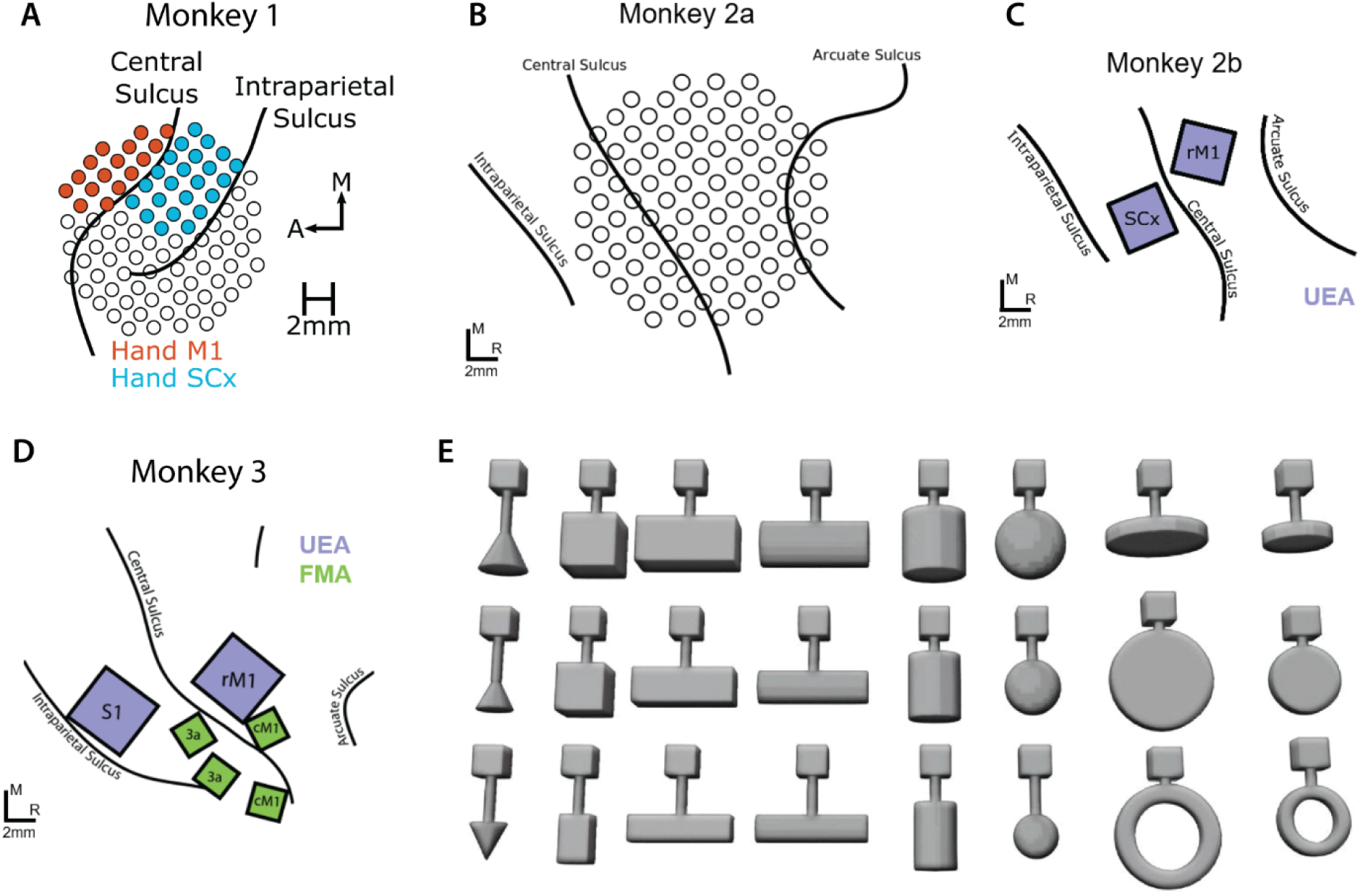
Electrode array placement locations and grasped objects. **ABCD**| In monkeys 1 (A) and 2a (B) the recordings were performed using semi-chronic Microdrive electrode arrays. In monkeys 2b (C) and 3 (D) the recordings were performed using Utah electrode arrays (purple) and floating microelectrode arrays (green). **E**| Objects used in the study. Subplots ABCD reproduced with permission from^10^. Subplot E reproduced with permission from^53^.

**Supplementary Figure 2.**
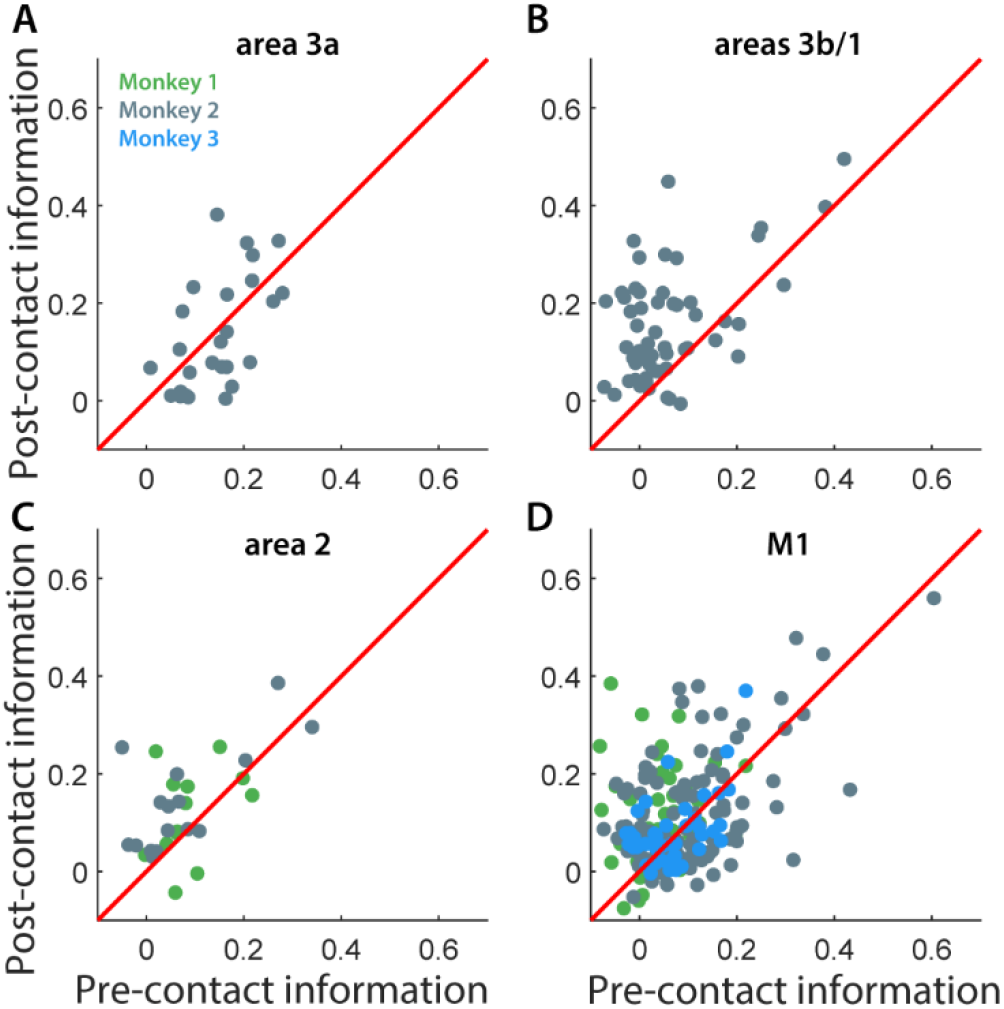
Object-specific signals carried by individual neurons in somatosensory areas. We computed the mutual information between the activity of each neuron and the identity of the grasped object for 400 ms windows before and after object contact period. We have subtracted shuffled mutual information for each neuron. High mutual information before contact would indicate primarily object-specific kinematic signals during preshaping of the hand in sensory areas with an addition of planning in M1, and after contact – both hand posture and contact forces. **A**| Mutual information between the activity of individual neurons in area 3a and object identity over the whole epoch. Each point denotes the informativeness of one neuron. Wilcoxon sign-rank test for 3a for monkey 2: p<0.001, r=0.82. **B** | Same as A for areas 3b/1. Monkey 2: p<0.001, r=-0.77. **C**| Same as A for area 2. Monkey 1: p=0.2, r=-0.36; Monkey 2: p<0.001, r=-0.85. **D**| Same as A for M1. Monkey 1: M1: p=0.06, r=-0.27; Monkey 2: p=0.003, r=-0.27; Monkey 3: p=0.29, r=0.16. The red line denotes identity.

**Supplementary Figure 3.**
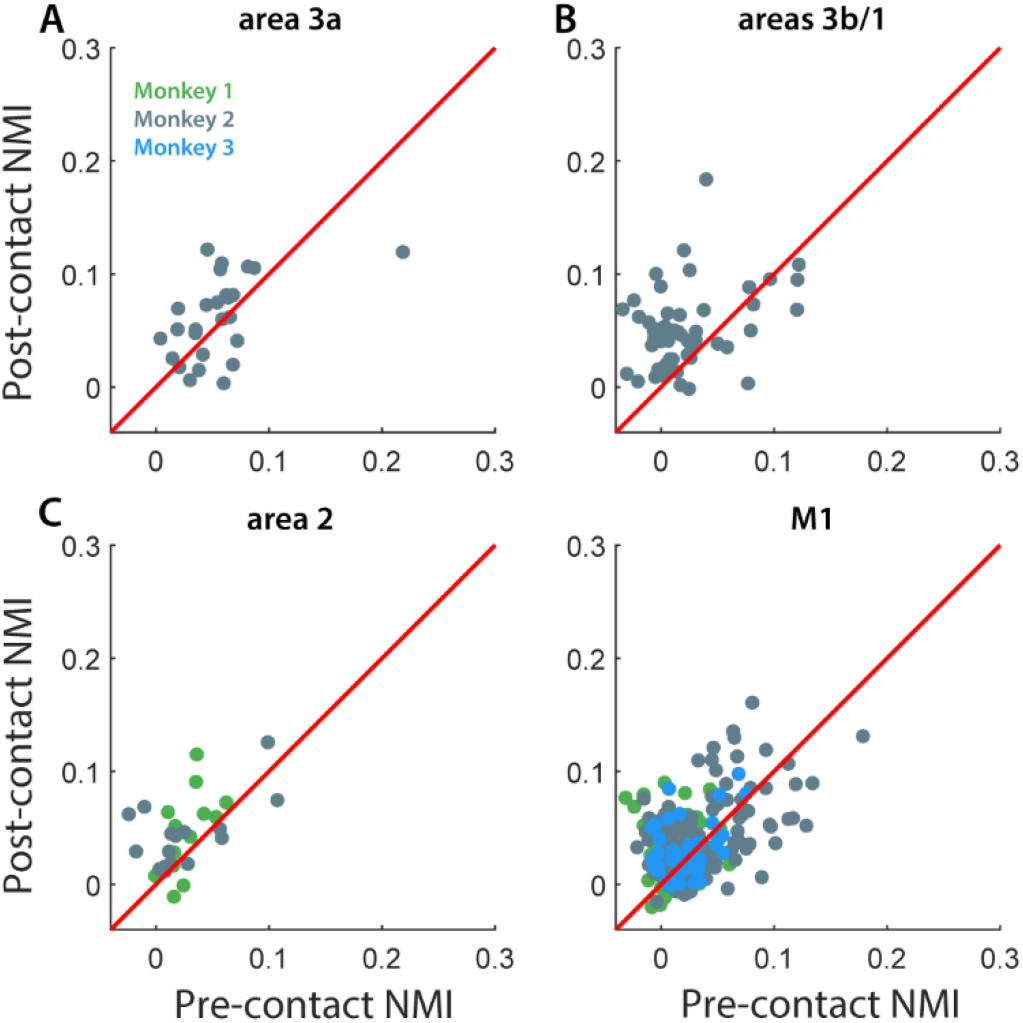
Normalized object-specific signals carried by individual neurons in somatosensory areas normalized to entropy of each neuron. We computed the normalized mutual information (NMI) between the activity of each neuron and the identity of the grasped object for 400 ms windows before and after object contact. We have subtracted shuffled NMI for each neuron. High NMI indicates consolidation of object-specific information. **A**| NMI between the activity of individual neurons in area 3a and object identity over the epoch. Each point denotes the informativeness of one neuron. Wilcoxon sign-rank test for 3a for monkey 2, respectively: p=0.73. **B** | Same as A for areas 3b/1. Monkey 2: p<0.001, r=-0.87. **C**| Same as A for area 2. Monkey 1: p=0.01, r=-0.71; Monkey 2: p<0.001, r=-0.87. **D**| Same as A for M1. Monkey 1: M1: p<0.001, r=-0.53; Monkey 2: p=0.003, r=-0.27; Monkey 3: p=0.29, r=0.16. The red line denotes identity.

**Supplementary Figure 4.**
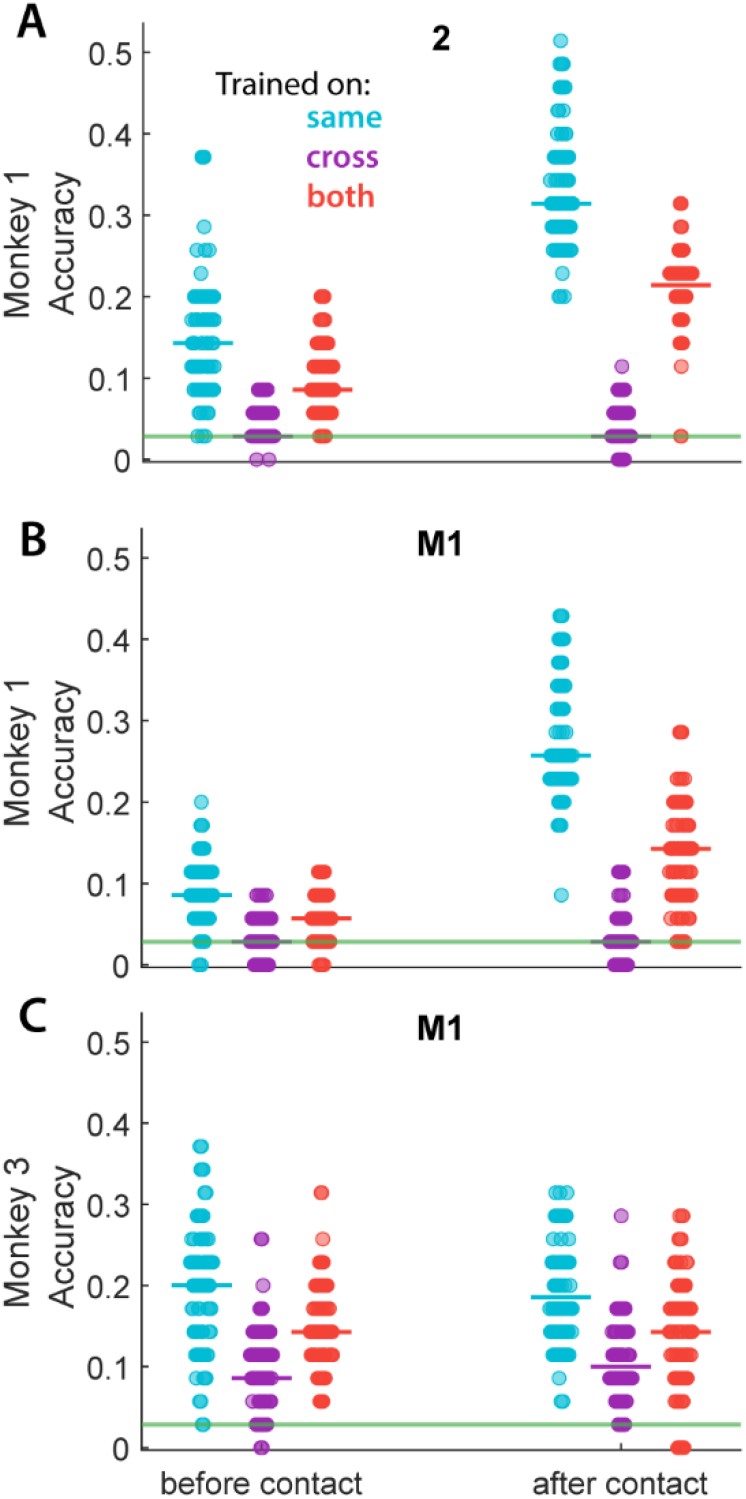
Stability of object-specific signals across epochs for monkeys 1 and 3. **A**| Performance of classifiers of object identity based on the responses of neurons in area 2 trained and tested within the same epoch (cyan), trained in one epoch and tested in the other (purple), and trained in both epochs then tested in each epoch separately (red). The “before contact” vs “after contact” column indicates the test epoch. Each point denotes the accuracy of classification based on a random allocation of training and testing data and the bold line denotes the median performance. Before and after same vs both: p<0.001, r=0.59 and r=0.95, respectively. **B**| Same as A for M1 of monkey 1. Before and after same vs both: p=0.06, r=0.15 and p<0.001, r=0.9, respectively. **C**| Same as A for M1 of monkey 3. Before and after same vs both: p<0.001, r=0.39 and p=0.02, r=0.2, respectively. Horizontal green line denotes chance (1/35).

**Supplementary Figure 5.**
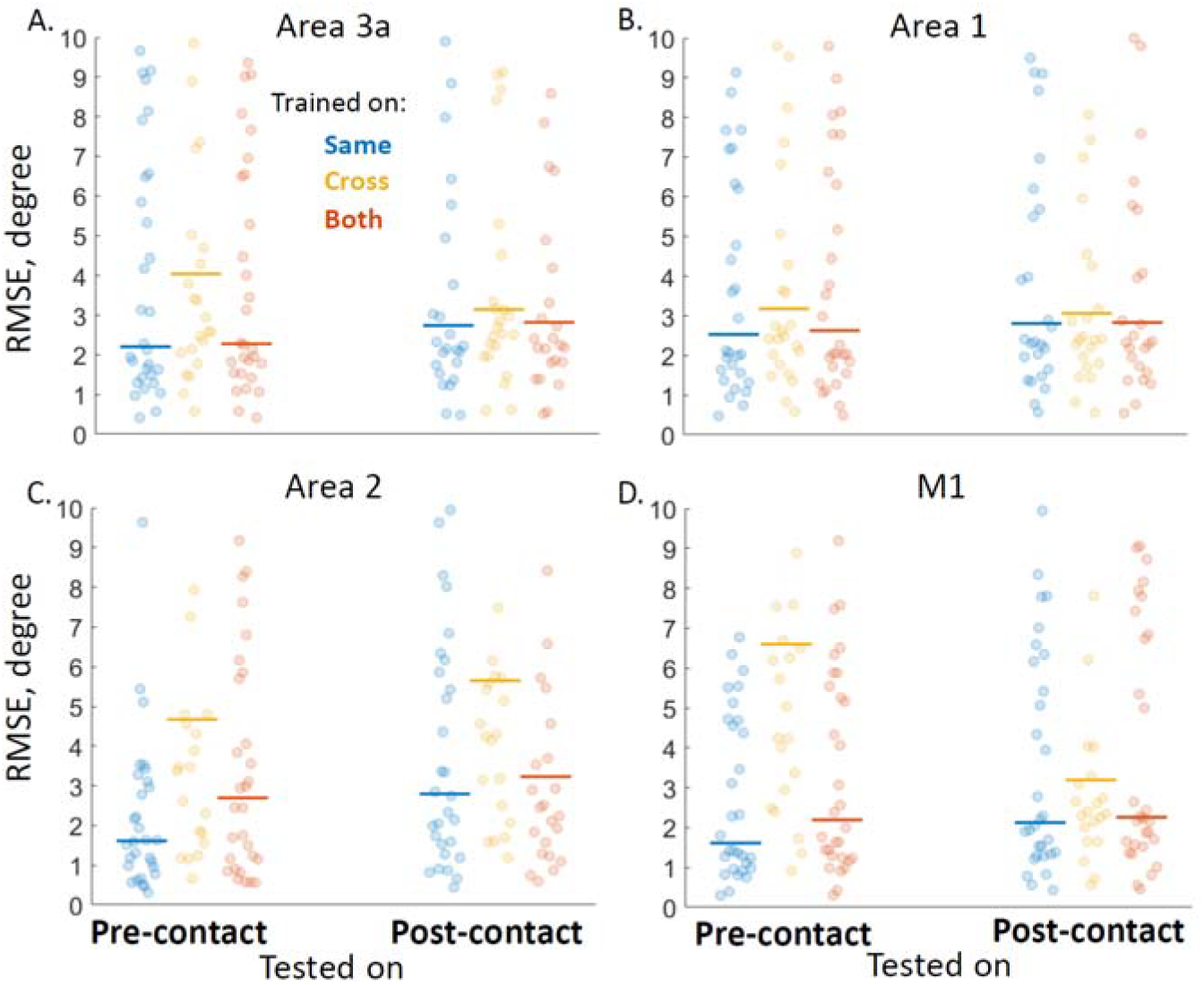
Stability of kinematic signal across epochs. A| Joint angle decoding performance using the spike counts of neurons from area 3a within a 200 ms window pre-contact (left) and post-contact (right). Each dot corresponds to one of 30 joint angles, and y-axis shows the root-mean-square error (RMSE) in degrees. B| Same as A for areas 3b/1. C| Same as A for area 2. D| Same as A for M1.

To directly investigate the similarity of the postural signal before and after contact, we decoded the hand joint angles from the pre-contact and post-contact neural responses. We found that joint angles can be predicted – within a few degree’s precision – from the neuronal spike counts. Marginally higher performance of the cross decoders in area 3a suggests that its representation is more consistent than M1’s. Area 2’s postural representations may be mixed with other signals and tend to vary more pre-contact and post-contact.

